# The E3 ubiquitin ligase TRIM9 regulates synaptic function and actin dynamics

**DOI:** 10.1101/2023.12.31.573790

**Authors:** Laura E. McCormick, Elliot B. Evans, Natalie K. Barker, Laura E. Herring, Graham H. Diering, Stephanie L. Gupton

**Affiliations:** Department of Cell Biology and Physiology, University of North Carolina at Chapel Hill, Chapel Hill, NC 27599; Michael Hooker Proteomics Core, University of North Carolina at Chapel Hill, Chapel Hill, NC 27599; Department of Pharmacology, University of North Carolina at Chapel Hill, Chapel Hill, NC 27599; Neuroscience Center, University of North Carolina at Chapel Hill, Chapel Hill, NC 27599; Lineberger Comprehensive Cancer Center, University of North Carolina at Chapel Hill, Chapel Hill, NC 27599

**Keywords:** filopodia, dendritic spine, synapse, TRIM9

## Abstract

During neuronal development, dynamic filopodia emerge from dendrites and mature into functional dendritic spines during synaptogenesis. Dendritic filopodia and spines respond to extracellular cues, influencing dendritic spine shape and size as well as synaptic function. Previously, the E3 ubiquitin ligase TRIM9 was shown to regulate filopodia in early stages of neuronal development, including netrin-1 dependent axon guidance and branching. Here we demonstrate TRIM9 also localizes to dendritic filopodia and spines of murine cortical and hippocampal neurons during synaptogenesis and is required for synaptic responses to netrin. In particular, TRIM9 is enriched in the post-synaptic density (PSD) within dendritic spines and loss of *Trim9* alters the PSD proteome, including the actin cytoskeleton landscape. While netrin exposure induces accumulation of the Arp2/3 complex and filamentous actin in dendritic spine heads, this response is disrupted by genetic deletion of *Trim9*. In addition, we document changes in the synaptic receptors associated with loss of *Trim9*. These defects converge on a loss of netrin-dependent increases in neuronal firing rates, indicating TRIM9 is required downstream of synaptic netrin-1 signaling. We propose TRIM9 regulates cytoskeletal dynamics in dendritic spines and is required for the proper response to synaptic stimuli.

## Introduction

During neuronal development, dynamic filopodia emerge from dendrites and mature into functional dendritic spines during synaptogenesis. Dendritic filopodia and spines respond to extracellular cues, influencing dendritic spine shape and size as well as synaptic function. Previously, the E3 ubiquitin ligase TRIM9 was shown to regulate filopodia in early stages of neuronal development, including netrin-1 dependent axon guidance and branching. Here we demonstrate TRIM9 also localizes to dendritic filopodia and spines of murine cortical and hippocampal neurons during synaptogenesis and is required for synaptic responses to netrin. In particular, TRIM9 is enriched in the post-synaptic density (PSD) within dendritic spines and loss of *Trim9* alters the PSD proteome, including the actin cytoskeleton landscape. While netrin exposure induces accumulation of the Arp2/3 complex and filamentous actin in dendritic spine heads, this response is disrupted by genetic deletion of *Trim9*. In addition, we document changes in the synaptic receptors associated with loss of *Trim9*. These defects converge on a loss of netrin-dependent increases in neuronal firing rates, indicating TRIM9 is required downstream of synaptic netrin-1 signaling. We propose TRIM9 regulates cytoskeletal dynamics in dendritic spines and is required for the proper response to synaptic stimuli.

E3 ubiquitin ligases are capable of ubiquitinating multiple substrates. Previously, we found TRIM9 is required for ubiquitination of two components of the filopodial tip complex, the actin polymerase VASP (Menon *et al*., 2015) and the netrin-1 receptor DCC (Plooster *et al*., 2017). The consequences of both ubiquitination events are consistent with a nondegradative modification that alters substrate function. Proximity-dependent biotin identification (BioID) in cultured neurons also identified a number of candidate binding partners and putative substrates involved in neuronal arborization and cytoskeletal organization (Menon *et al*., 2021). Although these experiments were completed before synaptogenesis began, we intriguingly identified several proteins related to synaptic structure as potential TRIM9 interactors.

Dendritic spines are hypothesized to mature from dendritic filopodia (Fiala *et al*., 1998). Although marked differences have been observed between growth cone filopodia and their dendritic counterparts (Korobova and Svitkina, 2010), a number of signaling pathways are conserved between both structures (McCormick and Gupton, 2020). Of note, the netrin-DCC pathway, classically studied for its role as an attractive axon guidance cue, was shown to promote dendritic filopodia formation, spine maturation, and synaptic plasticity (Goldman *et al*., 2013; Wong *et al*., 2019). Interestingly, netrin-1 induced activation of similar signaling cascades as occurs during long-term potentiation (synaptic strengthening), leading to an expansion of dendritic spine size and enhanced AMPA receptor insertion (Glasgow *et al*., 2018). As such, we hypothesize TRIM9 may localize to dendritic filopodia and regulate cytoskeletal changes involved in netrin-dependent dendritic spine morphogenesis.

Our previous in vivo studies suggested that TRIM9 may function at the synapse as well. We observed severe deficits in spatial learning and memory of *Trim9*^*-/-*^ mice in the Morris Water Maze test, compared to their littermate controls (Winkle *et al*., 2016). Furthermore, adult-born neurons in the dentate gyrus in *Trim9*^*-/-*^ mice also displayed altered dendritic architecture and a decreased number of dendritic spines (Winkle *et al*., 2016).

In this study, we investigate a synaptic role for TRIM9. We find that TRIM9 localizes to the tips of dendritic filopodia and spines and is enriched in the postsynaptic density (PSD). Loss of TRIM9 altered the proteome of the PSD in the forebrain, specifically the actin cytoskeleton landscape. Our in vitro results suggest that *Trim9*^*-/-*^ neurons also have altered synaptic receptor levels. Although the loss of *Trim9* does not alter baseline neuronal firing, we find that TRIM9 is required for appropriate response to netrin-1 treatment during synaptogenesis. Collectively, our results support an essential role for TRIM9 in appropriate synaptic signaling downstream of netrin-induced changes in dendritic spines.

## Results

### TRIM9 localizes to dendritic filopodia and spines during synaptogenesis

In early neuronal development, filopodia are required structures in neuritogenesis, axon branching, and axon guidance (Lebrand *et al*., 2004; Dent *et al*., 2007; Kwiatkowski *et al*., 2007). During later stages of development, filopodia also serve as dendritic spine precursors, probing the local environment to find synaptic partners. As TRIM9 regulates filopodial dynamics at early stages of neuronal development (Menon *et al*., 2015), we hypothesized that this ubiquitin ligase may also regulate filopodia and dendritic spine maturation during synaptogenesis. Interestingly, recent proteomic work in cultured murine neurons (DIV 21) identified TRIM9 as the 23^rd^ most abundant E3 ubiquitin ligase in whole cell lysates (Antico *et al*., 2023), suggesting TRIM9 is enriched in mature neurons and may function in dendritic spines as well.

To examine TRIM9 in more mature neurons, we transfected cultured hippocampal neurons with GFP-CAAX to mark the plasma membrane and visualize filopodia/spines and Myc-TRIM9 to evaluate protein localization at DIV 12 (**Fig 1A**). At this timepoint, dendritic filopodia have emerged and begun maturing into spines. Reminiscent of TRIM9 localization to growth cone filopodia tips, we observed TRIM9 at the tips of dendritic filopodia and within dendritic spines (**Fig 1A, Inset I and II**).

**Fig. 1.**
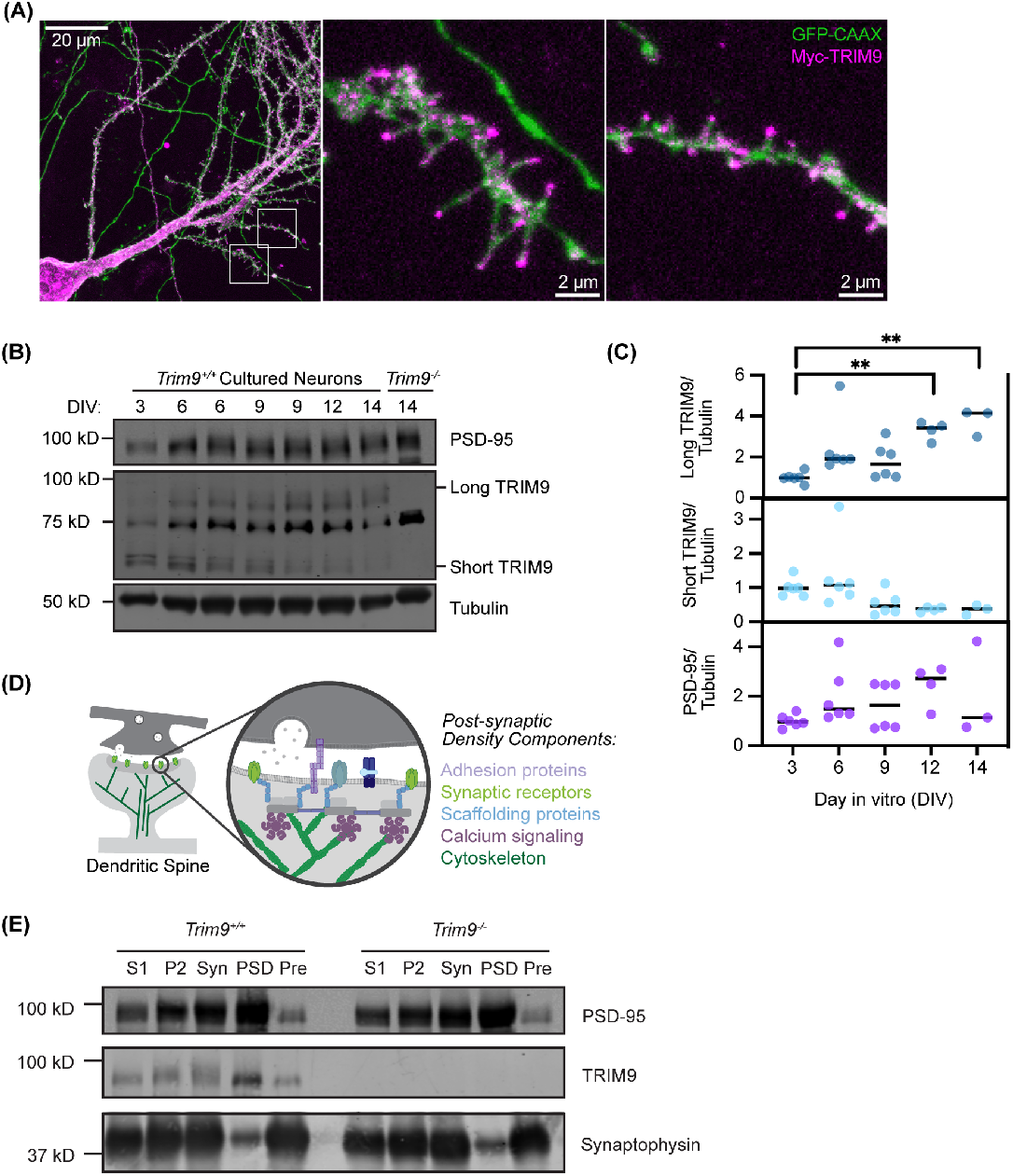
TRIM9 localizes to dendritic filopodia and spines and is enriched in the postsynaptic density during synaptogenesis. (A) Confocal microscopy (maximum projection image) of a fixed murine hippocampal neuron (DIV12) transfected with the membrane marker GFP-CAAX (green) and Myc-TRIM9 (magenta). Insets: Myc-TRIM9 localizes to the tips of dendritic filopodia and within dendritic spines. (B) Analysis of PSD-95, TRIM9 and βIII-tubulin levels by western blotting during in vitro neuronal development. The long TRIM9 isoform has an apparent molecular weight of 80 kDa, whereas the short isoform has an apparent molecular weight of 60 kDa. Both bands are missing in cultured *Trim9*^-/-^ neurons. The band at 75 kDa is a non-specific band recognized by the antibody that is also present in *Trim9*^*-/-*^ neurons. (C) Quantification of the long or short isoform of TRIM9 normalized to PSD-95 levels. Line indicates median for each condition. N = 3-6 technical replicates from 2-3 independent cultures. Kruskal-Wallis test with Dunn’s multiple comparisons. ** P < 0.01. (D) Schematic of the post-synaptic density (PSD), a protein-rich region within dendritic spines. (E) Representative immuneblots of post-synaptic density fractionation samples by western blot using the indicated antibodies. S1 (Supernatant 1), P2 (Pellet 2), Syn (Synaptosome), PSD (Postsynaptic density), Pre (Presynapse). 6 μg of protein was loaded per well.

To define how TRIM9 protein levels change during neuronal development, we cultured cortical murine neurons and examined TRIM9 protein levels during the first two weeks of in vitro growth **(Fig 1B)**. As previously reported (Winkle *et al*., 2016; Boyer *et al*., 2018), we observed multiple isoforms of TRIM9 through immunoblotting. We observed a single TRIM9 band migrating at approximately 80 kD, as well as a doublet migrating at 65-70 kD. Neither of these bands were present in cultures from *Trim9*^*-/-*^ neurons **(Fig 1B)**.

Like the well-characterized post-synaptic scaffolding protein PSD-95, we observed an increase in the abundance of the long isoform of TRIM9 as the culture matured in vitro. In contrast, the quantity of the lower molecular weight TRIM9 isoform decreased over time (**Fig 1C**). Of note, TRIM9 binds the netrin receptor DCC through its SPRY domain (Winkle *et al*., 2014)—a domain not present in the short TRIM9 isoform. Although netrin-1 was classically studied as an axon guidance cue (Boyer and Gupton, 2018), recent work demonstrated that netrin-1 also plays a role in synapse formation (Goldman *et al*., 2013). Collectively, these results indicate an isoform of TRIM9 involved in netrin signaling increases in abundance during synaptogenesis.

To examine the synaptic localization of TRIM9, we performed subcellular fractionation. The forebrains of juvenile mice (postnatal day 21, P21) were dissected and subjected to differential centrifugation to enrich for both the pre-synaptic fraction and the post-synaptic density (PSD), a dense conglomerate of proteins attached to the post-synaptic membrane. The PSD contains excitatory synaptic receptors, as well as scaffolding, signaling, and cytoskeletal proteins (**Fig 1D**). Similar to PSD-95, we observed a strong localization of TRIM9 in the PSD-enriched fraction compared to the early S1 fraction. In contrast, we observed an enrichment of the synaptic vesicle protein synaptophysin in the pre-synaptic enriched fraction (**Fig 1E)**.

### Loss of *Trim9* modifies the post-synaptic density proteome

To determine if the loss of *Trim9* altered the composition of the PSD, we performed quantitative mass spectrometry to examine the PSD proteome of *Trim9*^*+/+*^ and *Trim9*^*-/-*^ littermates (**Fig 2A**). Over 7,000 proteins were identified, and out of these, 109 proteins were changed in a statistically significant manner (p < 0.05) in the *Trim9*^*-/-*^ PSD (**Fig 2B, C**). Following Gene Ontology analysis, we observed the prominent enrichment of one pathway in our significantly different proteins: the actin cytoskeleton (**Fig 2D**). In particular, we observed decreases in six members of the Arp2/3 complex in the PSDs of *Trim9*^*-/-*^ mice (**Fig 2C**). To validate our proteomic results, we analyzed Arp2/3 protein levels in the PSD fractions by western blot by blotting for ArpC2, an obligate component of the complex. Interestingly, we observed a decrease in the level of subunit ArpC2 in male mice, but not female mice (**Fig 2E**).

**Fig. 2.**
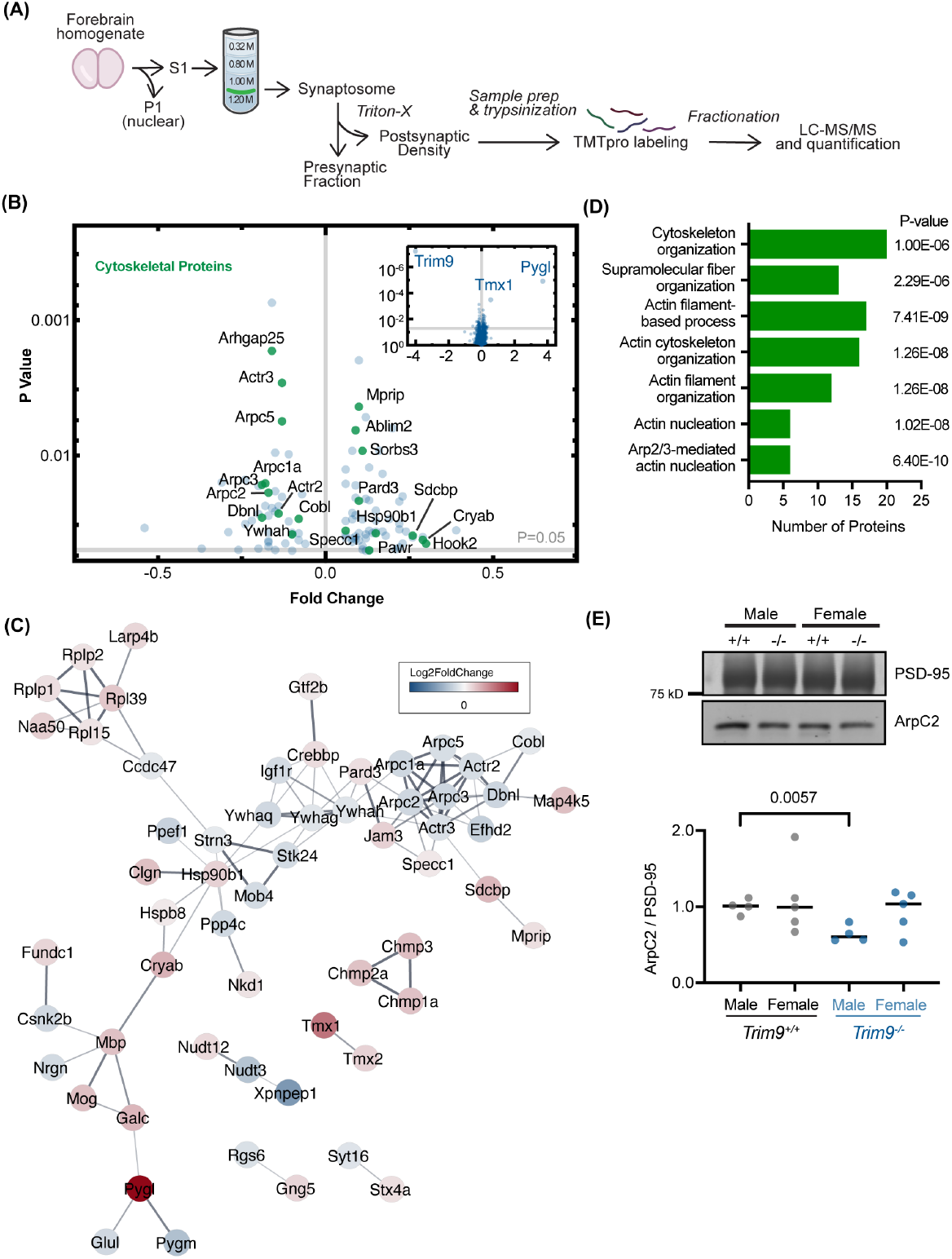
TRIM9 is enriched in the post-synaptic density (PSD) and loss of Trim9 alters the PSD proteome. (A) Schematic of subcellular fractionation of murine forebrains used to enrich the PSD fraction and subsequent preparative proteomic steps. (B) Volcano plot visualizing global proteomic changes in *Trim9*^*-/-*^ mice, highlighting significantly changed proteins (p < 0.05). Cytoskeletal proteins, identified by Gene Ontology (GO) analysis, are highlighted in green. Inset: Volcano plot with expanded x and y axis. (C) STRING Analysis highlighting known protein-protein interactions among the 109 proteins that were significantly changed in *Trim9*^*-/-*^ mice. The color of each protein node corresponds to the log2fold change of the protein. (D) GO analysis by biological processes. (E) A representative western blot and quantification of ArpC2 protein levels normalized to PSD-95 levels in the PSD enriched fraction. N=4-5 mice/genotype and sex. A Brown and Forsythe and Welch ANOVA test was completed with Dunnett’s T3 multiple comparisons.

As the actin cytoskeleton regulates the emergence and expansion of dendritic spines, we hypothesized that *Trim9*^*-/-*^ neurons may have altered dendritic spine number or shape. We cultured hippocampal neurons from *Trim9*^*+/+*^ and *Trim9*^*-/-*^ mice and quantified dendritic filopodia and spine number at DIV 12 (**Fig 3A-B**). At baseline we did not observe a significant difference in spine density between *Trim9*^*+/+*^ and *Trim9*^*-/-*^ neurons (**Fig 3B)**. Neurons were treated with sham media or netrin-1, previously shown to enhance dendritic filopodia formation, maturation, and synaptic transmission (Goldman *et al*., 2013; Glasgow *et al*., 2018) for four hours. We observed a 1.22-fold increase in the mean dendritic filopodia/spine number in *Trim9*^*+/+*^ neurons following netrin treatment (**Fig 3B**), consistent with previous work that netrin-1 stimulates filopodia formation and spine stabilization (Goldman *et al*., 2013). However, we observed no significant increase in filopodia/spine number following netrin treatment in *Trim9*^*-/-*^ neurons (**Fig 3B**), consistent with the requirement for TRIM9 in the netrin response observed at earlier stages of neuronal development.

**Fig. 3.**
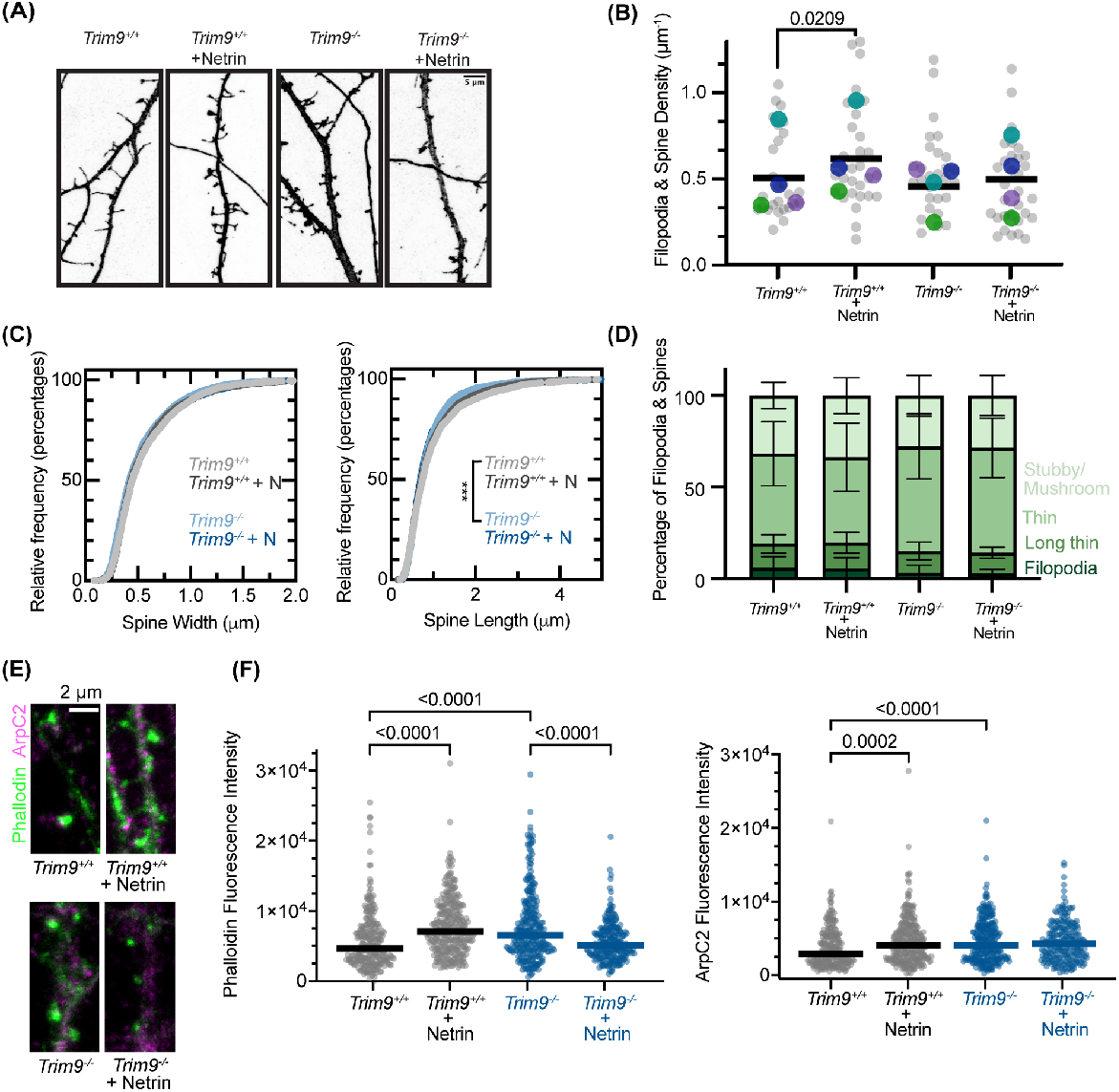
TRIM9 is required for netrin-dependent increases in filopodia/spine formation. (A) Confocal microscopy (maximum projection images) of branches from the primary dendrites of *Trim9*^*+/+*^ and *Trim9*^*-/-*^ hippocampal neurons (DIV12). Neurons were transfected with GFP-CAAX to visualize the plasma membrane. Cells were treated with netrin-1 (200 μg/mL) or sham media for four hours before fixation. (B) Quantification of dendritic filopodia and spine density (per μm of dendrite). >100 μm of dendrite was measured for each cell. N= 4 independent experiments encompassing 27-32 neurons in total (grey data points). The median for each experiment is visualized as the larger, colorful data points. RM one-way ANOVA with the Holm-Sidak’s multiple comparisons test. Line represents mean across all four experiments. (C) Relative frequency plots measuring dendritic spine length and width. 2493-3444 spines were measured for each condition across four independent experiments. Spine length: *Trim9*^*+/+*^ v. *Trim9*^*-/-*^, p < 0.0001 (Kolmogorov-Smirnov test). (D) Classification of dendritic spine subtypes based on width and length measurements. Categories were assigned as following: Filopodia (Length 2-5 μm, Width < 0.6 μm); Long thin (Length 1-2 μm, Width < 0.6 μm); Thin (Length < 1 μm, Width < 0.6 μm); Stubby/mushroom (Width > 0.6 μm).(E) Confocal imaging (maximum projection) of dendrites from cultured hippocampal neurons (DIV12) stained for phalloidin (green) and ArpC2, a component of the Arp2/3 complex (magenta). Quantification of phalloidin and ArpC2 levels in stubby and mushroom dendritic spines. N = 234-296 spines from 19-24 cells across three independent experiments. A Kruskal-Wallis test with Dunn’s multiple comparisons was completed. Line represents median for each condition.

As synaptic activity increases, the dendritic spine shortens in length and the spine head increases in diameter. Upon maturity, dendritic spines possess a characteristic mushroom-like shape. As such, we measured the length and width of each filopodia/spine. Netrin-1 did not alter spine width or length in *Trim9*^*+/+*^ or *Trim9*^*-/-*^ neurons (**Fig 3C**). However, *Trim9*^*-/-*^ neurons exhibited a small but significant decrease in dendritic spine length. As visual classification of spine morphology can be subjective, we used these width and length measurements spines to categorize spine sub-type in order of increasing maturity: filopodia; long thin; thin; mushroom/stubby. However, we did not observe changes in spine sub-type distribution across genotypes or netrin-1 treatment (**Fig 3D**).

To connect the cytoskeletal changes observed in the *Trim9*^*-/-*^ PSD proteomics with the altered dendritic spine phenotypes observed in vitro, we examined levels of the Arp2/3 complex and filamentous actin in the dendritic spines of cultured hippocampal neurons. We quantified the intensity of ArpC2, a component of the Arp2/3 complex, and phalloidin staining of filamentous actin in stubby and mushroom-shaped spines (**Fig 3E**). We observed a significant increase in filamentous actin in dendritic spines in *Trim9*^*+/+*^ neurons following four hours of netrin treatment, indicating an increase in polymerized actin in these structures. We also observed an increase in the level of ArpC2 following netrin-1 treatment in *Trim9*^*+/+*^ neurons. Interestingly, we observed elevated levels of both phalloidin and ArpC2 in dendritic spines of *Trim9*^*-/-*^ neurons at baseline, although a decrease was observed in phalloidin following netrin-1 treatment (**Fig 3E,F**). These results indicate loss of *Trim9* alters levels of postsynaptic ArpC2 and polymerized actin in vitro. Furthermore, this confirms TRIM9 is required for dendritic spine responses to netrin.

### Loss of *Trim9* alters synaptic receptor levels

Previous work suggested that netrin-1—through the transmembrane receptor DCC—leads to post-synaptic activation of the kinases PKA and PKC. In turn, this kinase activity results in the phosphorylation and insertion of the excitatory AMPAR GluA1 in the membrane (Glasgow *et al*., 2018, 2021). Furthermore, netrin-1 treatment enhances actin polymerization in dendritic spines through Src family kinase activation (Goldman *et al*., 2013). To probe the mechanisms underlying the failure of *Trim9*^*-/-*^ neurons to respond to netrin treatment, we utilized quantitative western blotting to examine the components of these netrin signaling pathways. *Trim9*^*+/+*^ and *Trim9*^*-/-*^ cortical neurons (DIV 12) were treated with netrin-1 for 45 minutes. We performed surface biotinylation to label proteins on the extracellular face of the plasma membrane and analyzed protein enrichment following streptavidin pulldown by western blotting. Although we did not detect significant changes in the surface level of excitatory GluA1, we observed an increase in the surface level of the inhibitory GABA_A_Rα-1 in *Trim9*^*-/-*^ neurons compared to *Trim9*^*+/+*^ controls (**Fig 4A,B**).

**Fig. 4.**
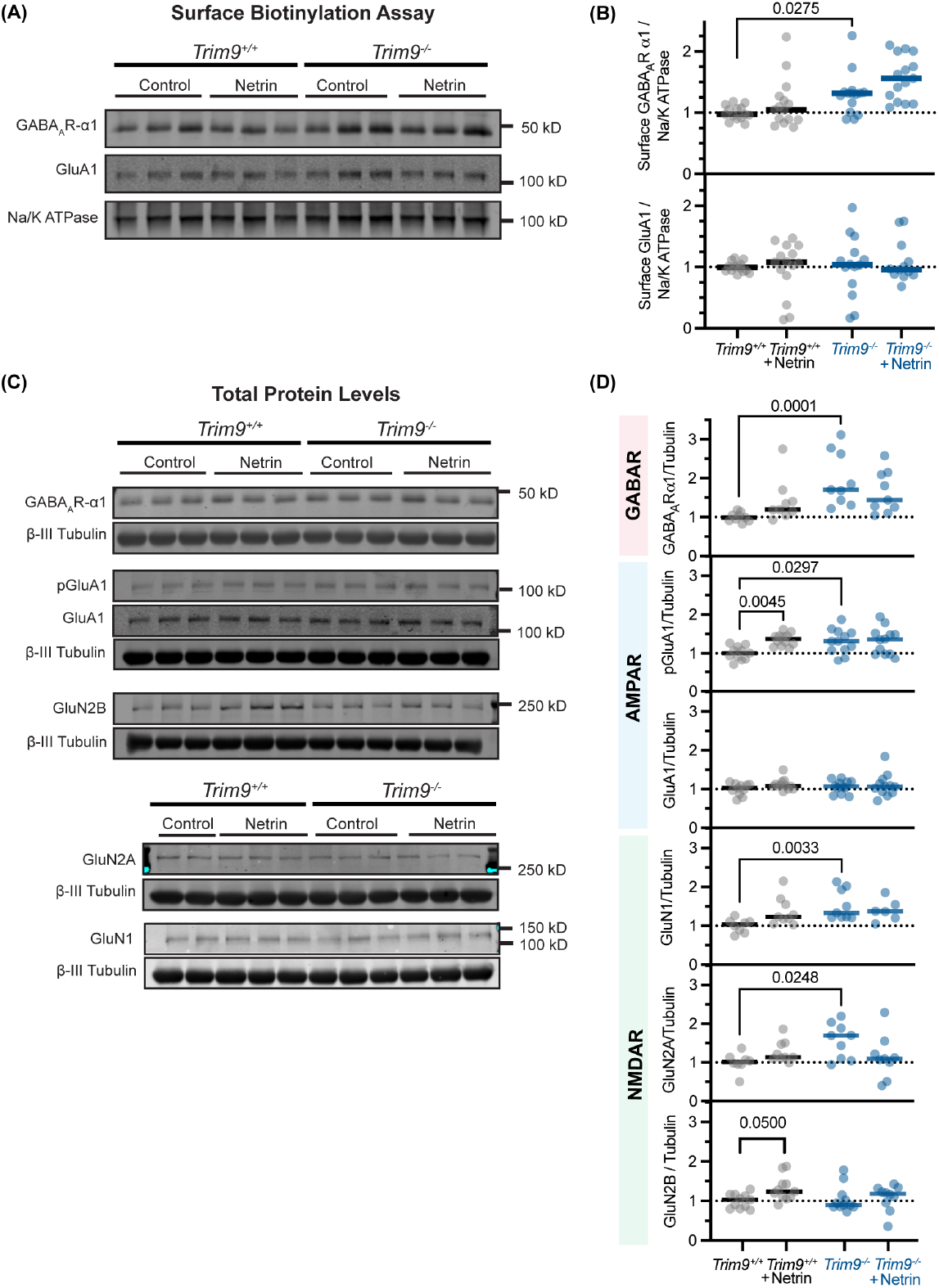
Loss of *Trim9* alters synaptic protein receptor levels. (A) Cultured cortical neurons (DIV14) were treated with netrin-1 (200 μg/mL) or sham media for forty-five minutes. Proteins at the cell surface were labeled with Sulpho-NHS-SS-Biotin followed by cell lysis and streptavidin pulldown. (B) Surface protein levels were analyzed by quantitative western blotting and normalized to Na/K ATPase. N = 11-12 technical replicates across 4 independent experiments (GABA_A_Rα-1). N = 9-12 technical replicates across 4 independent experiments (GluA1). A Kruskal-Wallis test with Dunn’s multiple comparisons was completed for each protein. Line represents median for each condition. (C) Western blotting of total cell lysate protein levels from biotinylation experiments. (D) Total protein levels were normalized to B-III tubulin. N = 9 technical replicates across 3 independent experiment for (GABA_A_Rα-1). N = 11-12 technical replicates across 4 independent experiments (pGluA1 and GluA1). N = 6-9 technical replicates across 2-3 independent experiment (GluN1). N = 8-9 technical replicates across 3 independent experiments (GluN2A). N = 7-9 technical replicates across 4 independent experiments (GluN2B). A Kruskal-Wallis test with Dunn’s multiple comparisons was completed for each protein. Line represents median for each condition.

To complement this quantification of receptors at the neuronal surface, protein levels in whole cell lysate were analyzed. Consistent with the biotinylation assay results, we observed an increase in the level of GABA_A_Rα-1 in *Trim9*^*-/-*^ neurons at baseline. Although the levels of AMPA receptors GluA1 (**Fig 4C,D**) and GluA2 (**Fig S2**) did not change following netrin-1 treatment in *Trim9*^*+/+*^ neurons, we did observe an increase in the amount of pGluA1 (S845). Interestingly, the baseline level of pGluA1 (S845) was elevated in *Trim9*^*-/-*^ neurons and did not further increase in the presence of netrin-1 (**Fig 4C, D**). pGluA1 is a modified form of the protein appreciated to enhance ion channel function, as well as resist endocytosis from the plasma membrane or enhance recycling (Diering and Huganir, 2018). Although we did not detect significant changes in surface GluA1 in the streptavidin pulldown, this increase in total pGluA1 suggests that the membrane retention of GluA1 may be increased in the absence of TRIM9.

Lastly, we also observed elevated levels of two NMDA receptors, GluN1 and GluN2A, in *Trim9*^*-/-*^ neurons. Although electrophysiology experiments suggested NMDAR activity was not altered downstream of netrin-DCC signaling, NMDAR activation by chemical LTP enhanced netrin-1 secretion in cultured hippocampal neurons (Glasgow *et al*., 2018). In contrast, GluN2B levels were not significantly changed in *Trim9*^*-/-*^ neurons, but we did observe an increase in GluN2B following netrin-1 treatment only in *Trim9*^*+/+*^ neurons (**Fig 4C,D**). We also did not detect netrin-1 or TRIM9-dependent changes in the levels of CaMKII, pCaMKII, gephyrin, B-actin, ArpC2 or Na/K ATPase in neuronal lysates (**Fig S2**). Collectively, loss of *Trim9* led to an increased level of pGluA1, mirroring the protein changes downstream of netrin-1 treatment in *Trim9*^*+/+*^ neurons. However, we also detected distinct protein level changes unique to *Trim9*^*-/-*^ neurons, including an increase in GluN1, GluN2A, and GABA_A_Rα-1.

### Loss of *Trim9* changes synaptic marker size and intensity

As we observed an increase in surface and total GABA_A_Rα-1 levels, we hypothesized that there may be a change in the number or composition of inhibitory synapses in *Trim9*^*-/-*^ neurons (DIV 12). To evaluate the ratio of inhibitory and excitatory synapses, we immunostained cultured hippocampal neurons with antibodies to both presynaptic and postsynaptic proteins. We utilized synaptophysin for a pan presynaptic marker, PSD-95 to mark excitatory post-synapses, and gephyrin to identify inhibitory postsynapses (**Fig 5A**). We identified colocalized PSD-95 and synaptophysin puncta between genotypes to mark excitatory synapses, as well as colocalized gephyrin, and synaptophysin puncta to denote inhibitory synapses. With these values, we calculated the ratio of excitatory to inhibitory synapses in the cultures and found no significant changes between *Trim9*^*+/+*^ and *Trim9*^*-/-*^ cultured neurons at DIV 12 (**Fig 5B**).

**Fig. 5.**
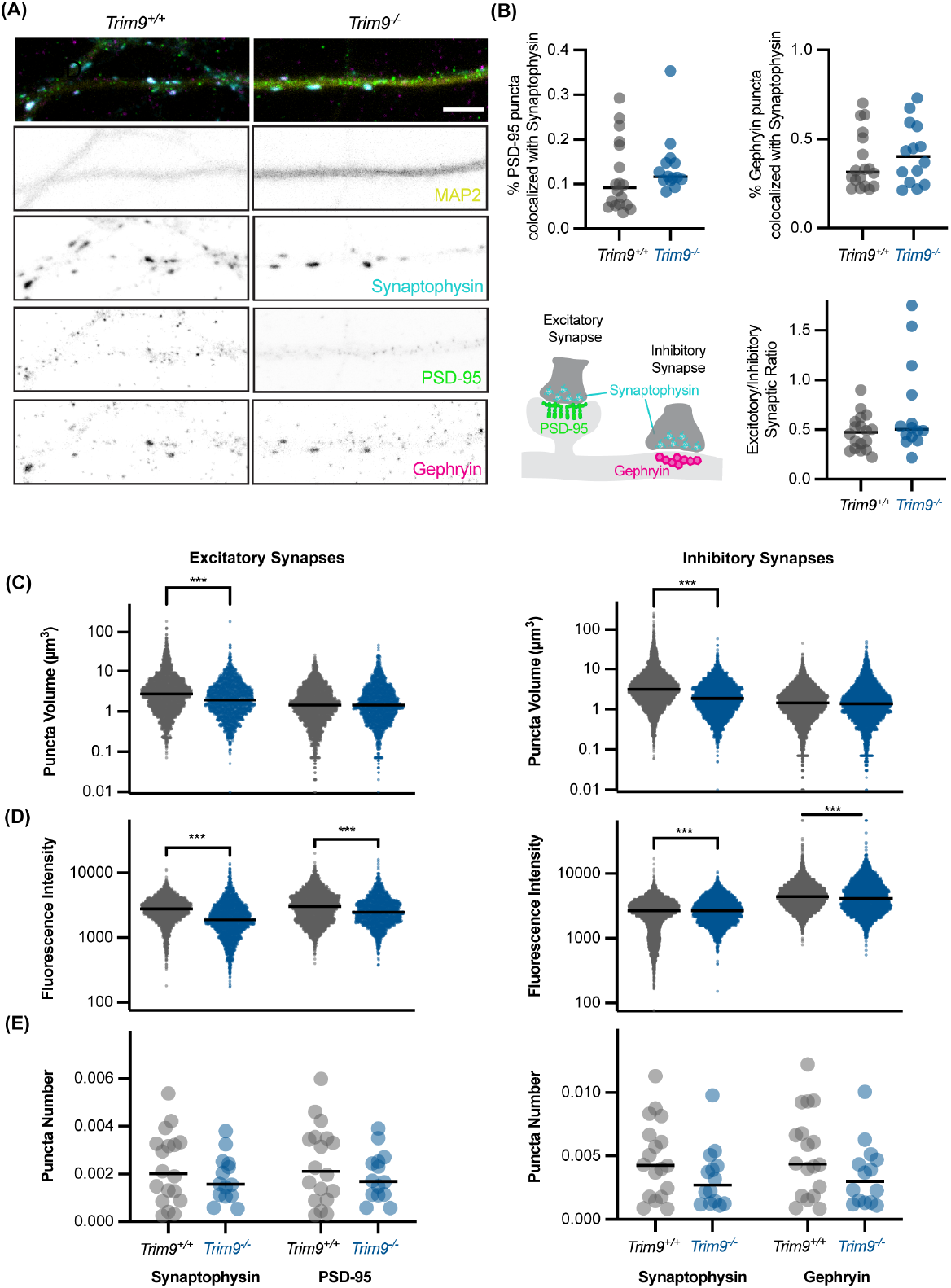
*Trim9*^*-/-*^ neurons do not exhibit an altered excitatory/inhibitory synapse ratio. (A) Confocal maximum projection of cultured hippocampal neurons (DIV12) stained with MAP2, as well the synaptophysin, PSD-95, and gephyrin. (B) Quantification of synaptophysin, PSD-95, and gephyrin puncta. Excitatory or inhibitory synapses per image were identified by detecting colocalized PSD-95 and synaptophysin puncta or gephyrin and synaptophysin puncta, respectively. Line represents median for each condition. N = 18 (*Trim9*^+/+^) and 14 (*Trim9*^*-/-*^) field of views across 3 independent experiments. (C) Puncta volume of colocalized synaptic puncta. Kruskal-Wallis test with Dunn’s multiple comparisons. N = 1872-7076 puncta from 18 (*Trim9*^*+/+*^) and 14 (*Trim9*^*-/-*^) field of views across 3 independent experiments. *** P < 0.001. (D) Mean fluorescence intensity measurements of colocalized synaptic puncta. Kruskal-Wallis test with Dunn’s multiple comparisons. N = 1872-7076 puncta from 18 (*Trim9*^*+/+*^) and 14 (*Trim9*^*-/-*^) field of views across 3 independent experiments. *** P < 0.001. (E) Total number of colocalized synaptic puncta per field of view normalized to MAP2 cumulative volume. N = 18 (*Trim9*^*+/+*^) and 14 (*Trim9*^*-/-*^) field of views across 3 independent experiments.

Next, we characterized the size and fluorescence intensity of the colocalized pre- and post-synaptic puncta. At excitatory synapses, we observed a significant decrease in the volume and fluorescence intensity of synaptophysin puncta in *Trim9*^*-/-*^ neurons, as well as the fluorescence intensity of PSD-95 puncta (**Fig 5C,D)**. These observations were consistent at inhibitory synapses; In *Trim9*^*-/-*^ neurons, we observed a significant decrease in the volume and fluorescence intensity of synaptophysin and a decrease in the fluorescence intensity of gephyrin puncta (**Fig 5C,D**). We did not detect significant changes in the overall number of colocalized synaptic puncta (**Fig 5E**). Collectively, these results suggest there is not a change in the number of excitatory/inhibitory synapses in *Trim9*^*-/-*^ neurons, but there may be a decrease in synaptic strength.

### TRIM9 is required for netrin-dependent increases in synaptic activity

To determine if TRIM9 plays a role in synaptic function, we utilized microelectrode arrays to measure the electrical activity of cortical neurons. In this assay, neurons are cultured on specialized plates containing electrodes (**Fig 6A**). Neuronal firing was recorded during synapse formation and maturation at DIV 8, 10, 12, 13, and 14. We did not detect baseline differences in the mean firing rate of *Trim9*^*+/+*^ and *Trim9*^*-/-*^neurons at these timepoints (**Fig 6B,C**). Neurons were also treated with sham media or netrin-1 at DIV 13. Consistent with previous results demonstrating netrin-1 bath perfusion enhanced AMPA receptor-mediated eEPSCs in hippocampal slices (Glasgow *et al*., 2018), we observed an increase in the mean firing rate of *Trim9*^*+/+*^ neurons following netrin-1 treatment over the course of six hours. In contrast, the mean firing rate of *Trim9*^*-/-*^ neurons was unchanged following netrin-1 treatment (**Fig 6D**), suggesting that TRIM9 is required for synaptic responses to netrin and downstream synaptic netrin-1 signaling.

**Fig. 6.**
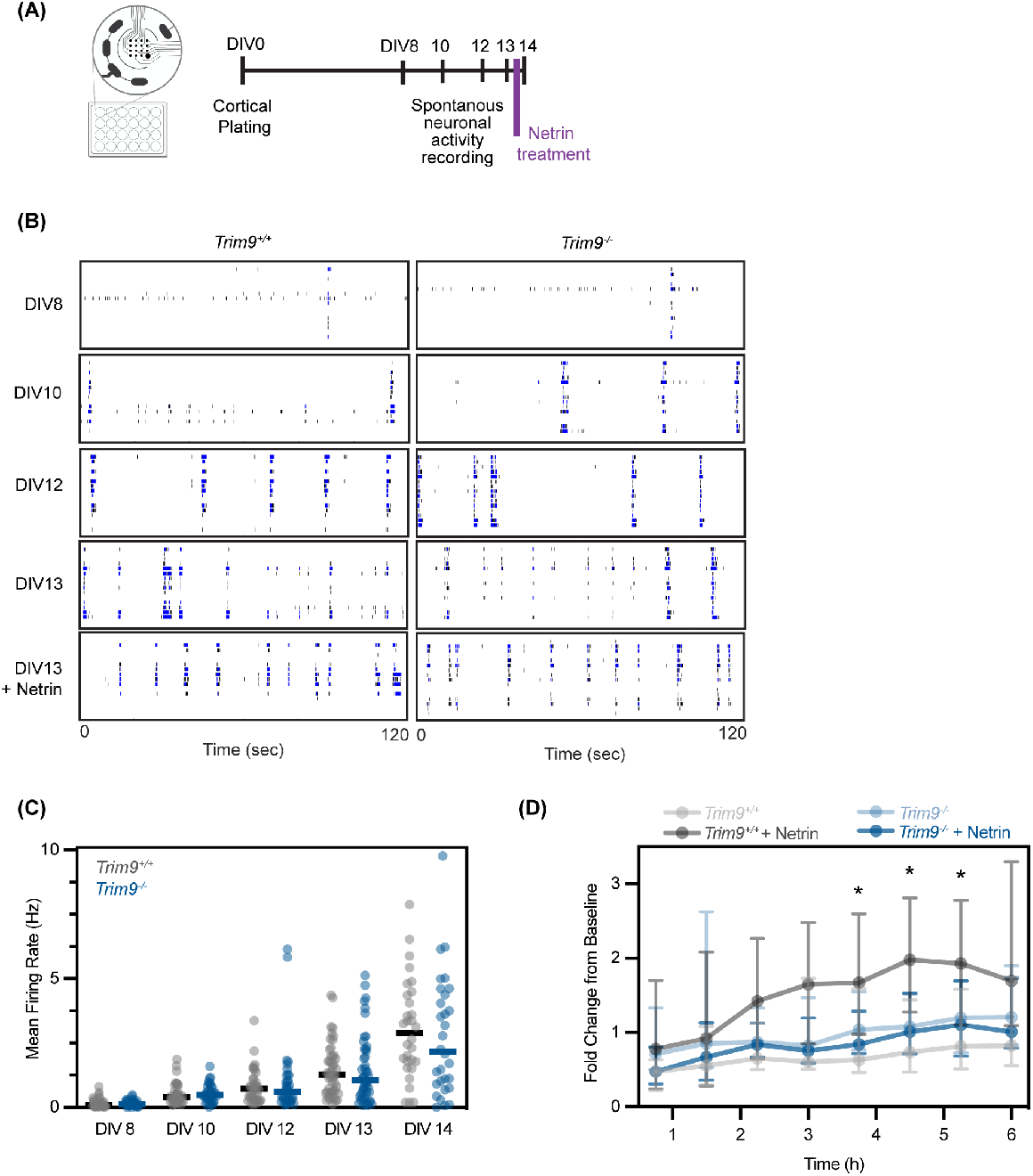
TRIM9 is required for netrin-dependent increases in neuronal firing. (A) Microelectrode array schematic. Neurons were plated on specialized plates containing electrodes and activity was measured at DIV 8, 10, 12, and 13 before netrin-1 treatment. (B) Representative raster plots of *Trim9*^*+/+*^ and *Trim9*^*-/-*^ cortical cultures from DIV 8-DIV 13. (C) Mean firing rate measurements of DIV 8-14 neurons. N = 30-48 replicates from 3-4 experiments. Line denotes median for each condition. (D) Change in the mean firing rate after sham media or netrin-1 (200 μg/mL) treatment over six hours at DIV13. Median firing rate with interquartile range plotted. N = 24 replicates from 4 independent experiments. * P < 0.05. Two-way ANOVA with Tukey’s multiple comparison test.

## Discussion

Here we show the brain-enriched E3 ubiquitin ligase TRIM9 localizes to dendritic spines and filopodia during synaptogenesis in vitro. Furthermore, TRIM9 localizes to the PSD within dendritic spines in the forebrain at P21, a period of synaptogenesis. Global PSD proteomics found protein level changes in numerous proteins that regulate the cytoskeleton, suggesting that the actin architecture in *Trim9*^*-/-*^ dendritic spines may be altered. Through both biochemical and microscopy-based assays, we demonstrated that TRIM9 was required for netrin-1-dependent changes in dendritic spine density and spine composition. Furthermore, we observed elevated levels of both excitatory (GluN1, GluN2, pGluA1) and inhibitory (GABA_A_Rα-1) receptors in cultured *Trim9*^*-/-*^ neurons during synaptogenesis. Despite these increased levels of receptors, we observed decreased volume and fluorescence intensity of pre- and post-synaptic marker puncta in *Trim9*^*-/-*^ neurons via immunofluorescence. Lastly, we demonstrate TRIM9 is required for netrin-1 dependent changes in neuronal firing.

### TRIM9 expression during development

Our previous work demonstrated that TRIM9 is expressed in both the embryonic and adult murine brain (Winkle *et al*., 2014; Boyer *et al*., 2018). In the developing mouse cortex, TRIM9 protein is detected during late embryonic stages, and subsequently increases in abundance, with levels peaking during the first two postnatal weeks and coinciding with periods of rapid synaptogenesis. Although expression levels decrease over time, TRIM9 persists in the adult cortex (Boyer *et al*., 2018).

Multiple isoforms of murine TRIM9 are predicted from cDNA and we find several TRIM9 reactive bands of different molecular weight by SDS-PAGE and immunoblotting that are absent in the *Trim9*^*-/-*^ neurons that likely correspond to the long and short isoforms of TRIM9 that differ at the 3’ end. Although the complete individual roles of each isoform have not been fully elucidated, the short isoform of TRIM9 (sTRIM9) was shown to auto-ubiquitinate and regulate the inflammatory response to viral infection in immortalized cell lines (Qin *et al*., 2016). sTRIM9 also regulates ubiquitination of the kinase MKK6 in glioblastoma progression (Liu *et al*., 2018). Of note, the short isoform also lacks the SPRY domain, which we previously showed directly interacts with the netrin receptor DCC (Winkle *et al*., 2014). We found this isoform declined in abundance as the neuronal cultures aged (**Fig 1B, C**). The prevalence of the long TRIM9 isoform and the failure of *Trim9*^*-/-*^ neurons to appropriately respond to netrin-1 suggests these proteins interact at the synapse.

### The actin cytoskeleton regulates synaptic receptor trafficking and organization

We observe numerous synaptic protein level changes in cultured *Trim9*^*-/-*^ neurons. In whole cell lysates, we detect significant increases in both excitatory (GluN1, GluN2, pGluA1) and inhibitory (GABA_A_Rα-1) synaptic receptors (**Fig 4C,D**). Likewise, polymerized actin and Arp2/3 levels in *Trim9*^*-/-*^ dendritic spines were elevated (as quantified by immunofluorescence, **Fig 2E,F**). However, we surprisingly demonstrated a decrease in the volume and fluorescence intensity of pre- and post-synaptic puncta in cultured *Trim9*^*-/-*^ neurons (**Fig 5C,D**). These bidirectional changes in synaptic organization are intriguing and suggest TRIM9-dependent changes in actin architecture may alter retention of receptors at the synapse.

This relationship between actin polymerization and receptor dynamics is well studied. At the excitatory post-synapse, the actin cytoskeleton regulates AMPA receptor trafficking (Hanley, 2014) and physically limits receptor mobility at the membrane (Rust *et al*., 2010). Likewise, actin is hypothesized to tether NMDARs in place (Allison *et al*., 1998). Furthermore, the scaffolding protein PSD-95 is linked to polymerized actin by α-actinin. This binding event is required for PSD-95-mediated GluA1 accumulation at synapses (Matt *et al*., 2018). Although we observed increased levels of excitatory receptors in *Trim9*^*-/-*^ whole cell lysates, we did not detect significantly increased levels of GluA1 at the cell surface. We can hypothesize the decreased levels of PSD-95 observed by immunocytochemistry in *Trim9*^*-/-*^ synapses may prevent effective GluA1 retention at the membrane. In turn, the increased actin polymerization and augmented levels of receptors—including the activity-enhancing pGluA1 modification— may compensate for decreased post-synaptic scaffolding. However, determining the primary change occurring in the *Trim9*^*-/-*^ post-synapse versus downstream compensatory effects requires further experimentation. Interestingly, several experiments in cultured neurons demonstrated that the phenotypes of *Trim9*^*-/-*^ neurons mirrored those of *Trim9*^*+/+*^ neurons treated with netrin, including elevated phalloidin and ArpC2 staining in dendritic spines (**Fig 3E, F**) and pGluA1 levels in whole cell lysate (**Fig 4C,D**). This similarity may suggest the signaling pathways that occur downstream of netrin-1 are enhanced at baseline in *Trim9*^*-/-*^ neurons.

Although our work primarily focused on excitatory synapses, we also observed enhanced levels of inhibitory GABA_A_Rα-1 in both whole cell lysate and at the neuronal surface of *Trim9*^*-/-*^ neurons (**Fig 4A-D**). However, we did not observe statistically significant changes in the total number of inhibitory synapses or the excitatory/inhibitory synapse ratio in *Trim9*^*-/-*^ cultures (**Fig 5B, E**). Interestingly, recent work demonstrated GABA_A_Rα-1 binds secreted netrin-1. This interaction enhances conductance or flow of ions through GABA_A_Rα-1 and regulates homeostatic scaling (Chan *et al*., 2022). To understand the mechanism underlying this enhanced receptor level, future work must determine whether TRIM9 binds GABA_A_Rα-1 and/or participates in down-stream homeostatic signaling. Furthermore, experiments investigating the role of netrin-1 in DCC and GABA_A_Rα-1 signaling have been completed in isolation; whether these two receptors compete for netrin-1 binding or how these two pathways combinatorically impact neuronal activity is unknown.

### TRIM9 is required to properly respond to netrin-1

Despite changes in synaptic receptor levels detected by immunoblot (**Fig 4**) and changes in synaptic markers detected by immunofluorescence (**Fig 5C-F**), analysis of neuronal network firing dynamics using MEA did not reveal deficits during baseline culture conditions (**Fig 6C**). These data suggest that *Trim9*^*-/-*^ neurons must compensate for the loss of the E3 ligase at baseline. However, we observed that *Trim9*^*-/-*^ neurons were not able to respond to netrin-1 treatment in MEA assays (**Fig 6D**), suggesting compensation may not work at this timescale or in this signaling pathway. Although we cannot make conclusions on synapse-specific substrates of TRIM9, our proteomic results suggest that the altered cytoskeletal landscape of *Trim9*^*-/-*^ synapses may interfere with the rapid structural rearrangement needed for synaptic stimuli responses.

Future work is required to evaluate the response of *Trim9*^*-/-*^ neurons during synaptic plasticity or activity-dependent synaptic strengthening. As netrin-1 treatment is proposed to share similar pathways as structural plasticity (Glasgow *et al*., 2021), we may expect similar deficits in the ability of *Trim9*^*-/-*^ neurons to respond to plasticity inducing stimuli.

In particular, as pGluA1 is a required modification for the progression of synaptic strengthening (Diering and Huganir, 2018), we expect that the signaling pathway underlying this phosphorylation is already operating at capacity in the *Trim9*^*-/-*^ neurons. As such, we hypothesize *Trim9*^*-/-*^ neurons must compensate to avoid excitotoxicity from elevated signaling. Subsequent deficits in synaptic stimuli response may underly the dramatic spatial memory and learning phenotypes of *Trim9*^*-/-*^ mice (Winkle *et al*., 2016).

### Linking axon guidance and synaptogenesis

The signaling pathways downstream of netrin-1 in the growth cone are numerous and complex—including both actin and microtubule cytoskeletal remodeling, calcium signaling via PLCγ (Xie *et al*., 2006), and MAPK signaling (Forcet *et al*., 2002; Campbell and Holt, 2003). Our previous work has demonstrated TRIM9 is required for netrin-1 response in the growth cone. In particular, we have shown that TRIM9 directly interacts and ubiquitinates DCC (Plooster *et al*., 2017), as well as the actin polymerase VASP (Menon *et al*., 2015; Boyer *et al*., 2020). Although our lab has worked to identify new substrates and interacting partners of TRIM9 during axon guidance (Menon *et al*., 2020, 2021), there are still dozens of unvalidated candidates waiting to be explored.

Over the past decade, netrin-1 has been shown to influence signaling cascades in dendritic spines (Goldman *et al*., 2013; Horn *et al*., 2013; Glasgow *et al*., 2018, 2021; Wong *et al*., 2019). Although these synaptic cascades lead to the expansion of dendritic spines rather than expansion of the growth cone, both responses require remodeling of the actin cytoskeleton. Here we demonstrate that TRIM9 is required for the netrin response in more mature neurons, but the precise level at which this happens is still unclear. Several validated TRIM9 interaction partners originally identified in the growth cone—DCC, Myo16, VASP, PRG-1, and Kif1a—are known to localize to the synapse as well (Trimbuch *et al*., 2009; Lin *et al*., 2010; Goldman *et al*., 2013; Li *et al*., 2016; Roesler *et al*., 2019; Menon *et al*., 2021). However, netrin-1 also influences synapse specific signaling pathways, such as the regulation of AMPA receptor GluA1 (Glasgow *et al*., 2018). As such, we believe many exciting new substrates of TRIM9 are waiting to be identified at the synapse. While we have identified a subset of proteins that are changed in the PSDs of *Trim9*^*-/-*^ mice, additional studies are required to identify true TRIM9 substrates versus compensatory changes.

### Pre-synaptic v. post-synaptic contributions of TRIM9

While this work has focused on the role of TRIM9 in dendritic spines, we embrace the possibility that the ligase may function on both sides of the synapse. Although our murine forebrain fractionations suggest TRIM9 levels are elevated in the PSD-enriched fraction, TRIM9 was recently identified in a pre-synaptic proteomic study (O’Neil *et al*., 2021) and we detected TRIM9 protein in the pre-synapse in fractionation approaches as well (**Fig 1E**). Furthermore, we detected decreased volume and intensity of synaptophysin puncta in *Trim9*^*-/-*^ neurons (**Fig 5C,D**). Interestingly, TRIM9 was first discovered as a novel interaction partner of the t-SNARE protein SNAP25 in the rat brain and shown to interact with synaptic vesicles (Li *et al*., 2001). Our previous work confirmed that TRIM9 interacts with SNAP25. We additionally found that TRIM9 regulated constitutive exocytosis in developing neurons prior to synapse formation (Winkle *et al*., 2014). Collectively, these data suggest that TRIM9 may be a multidimensional synaptic regulator.

## Methods

### Data Availability

The proteomics dataset generated and analyzed in this study are available in the Proteomics Identification Database (PRIDE) repository under project identifier PXD048045.

### Plasmids

*In house*: Creation of Myc-TRIM9 full-length was previously described (Winkle et al., 2016). *Acquired*: GFP-hRas CAAX (Richard Cheney, University of North Carolina).

### Antibodies

- Rabbit polyclonal anti-p34-Arc/ARPC2 (Sigma 07-227-I-100UG): 1/300 IF; 1/1000 WB
- Mouse monoclonal β-actin (ProteinTech 66009-1-Ig): 1/5000-10000 WB
- Mouse monoclonal β-III Tubulin (Biolegend 801202): 1/5000 WB
- Rabbit monoclonal CaMKII (Cell Signaling 33625): 1/2500 WB
- Rabbit monoclonal pCaMKII (Thr286) (Cell Signaling D21E4): 1/1000 WB
- Rabbit polyclonal GABAAR-1 (Sigma Aldrich 06-868): 1/1000 WB
- Mouse monoclonal Gephyrin (Antibodies, Inc: 75-465): 1/200 WB
- Chicken polyclonal GFP (GFP-1010, Aves Labs): 1/500 IF
- Mouse monoclonal GluA1/GluR1 Glutamate receptor (DSHB - U Iowa N355/1-s): 0.36 μg/mL WB
- Rabbit polyclonal pGluA1 (pS845) (Millipore AB5849): 1/1000 WB
- Mouse monoclonal GluA2/GluR2 Glutamate receptor (DSHB - U Iowa L21/32-s): 0.44 μg/mL WB
- Rabbit polyclonal GluN1(Cell Signaling 5704): 1/1000 WB
- Rabbit polyclonal GluN2A (Cell Signaling 4205): 1/1000 WB
- Rabbit polyclonal GluN2B (Cell signaling 14544): 1/1000 WB
- Chicken polyclonal MAP2 (Fisher 01-670-262): 1/5000 IF
- Mouse monoclonal against c-Myc (9E10): 1/400 IF; 1/2000 WB
- Rabbit polyclonal Na/K ATPase (Abcam ab76020): 1/10000 WB
- Mouse monoclonal PSD-95 (NeuroMab 75-028): 1/500,000 WB
- Rabbit polyclonal PSD-95 (Abcam 18258): 1/500 IF
- Guinea pig polyclonal Synaptophysin (Synaptic Systems 101 004): 1/500 IF
- Rabbit polyclonal Synaptophysin (Proteintech 17785-1-AP): 1/2000 WB
- Rabbit polyclonal TRIM9 (generated in house against murine TRIM9 recombinant protein amino acids 158–271): 1/2000 WB

### Recombinant Netrin-1

Recombinant chicken myc-Netrin-1 was purified from HEK293 cells (Serafini et al., 1994; Lebrand et al., 2004). Neurons were treated with 200 μg/mL for 45 minutes – 6 hours depending on the experiment.

### Experimental Models

All mouse lines were on a C57BL/6J background and bred at the University of North Carolina with approval from the Institutional Animal Care and Use Committee. Creation of the *Trim9*^/^mouse line was previously described (Winkle *et al*., 2014). Timed pregnant females were obtained by placing male and female mice together overnight; the following day was denoted as E0.5 if the female had a vaginal plug.

### Neuronal Culture and Transfection

Hippocampal and cortical neuronal cultures were prepared from E17.5 C57Bl/6J mice. Tissue was dissected from HEPES/HBSS buffer, digested for 20 minutes with trypsin, washed with trypsin-quenching media (Neurobasal A supplemented with B27, 2 mM Glutamax, 5% FBS, and 1% penicillin/streptamycin) and dissociated with a p1000 pipette tip. Cells were then passed through at 70 μm cell strainer. Neurons were plated in trypsin quenching media onto PDL-coated coverslips or dishes. One to three hours after plating, the media was changed into serum free media (Neurobasal A media supplemented with B27 and 2 mM Glutamax). Three days after plating (DIV 3), cells were fed with an equivolume of glia conditioned media. At DIV 7, one-half of the media was removed and replaced with fresh glia conditioned media.

Neuronal transfections were performed as previously described (Plooster *et al*., 2021) based on a modified Lipofectamine2000 protocol. For each well of a 6 well plate, 6 μL of Lipofectamine2000 was mixed with 100 μL of Neurobasal media. 1.5-3 μg of DNA was also mixed with 100 μL of Neurobasal. The Lipofectamine and DNA were gently combined and incubated for 20 minutes at RT. Excess media from neurons was removed and saved—leaving 1 mL left in the well—and the Lipofectamine-DNA mixture was gently added dropwise to the neurons. After a 35 minute incubation at 37°C, all of the media was removed and replaced with a 50-50 combination of saved media and fresh media. Transfections were completed on DIV 7 before the scheduled feeding.

### Microelectrode Array

Microelectrode array assays were performed with a Maestro Edge (Axion Biosystems). Twenty-four well plates (Axion Biosystems) were coated overnight with 1 mg/mL PDL in borate buffer. 190,000 cortical neurons were seeded per well and fed at DIV 3 and DIV 7 as described above. Basal recordings were completed at DIV 8, 10, 12, and 14 for 10 minutes. Netrin treatments were completed with DIV 13 with 200 μg/mL of netrin-1 or sham media and recorded for 6 hours (a 5-minute reading was completed every 45 minutes). Data was analyzed with the Axion software.

### Immunostaining and Imaging

For fixed cell imaging experiments, #1.5 German glass coverslips (25 mm, Electron Microscopy Services #72290-12) were cleaned with a Harrick Plasma Cleaner (PDC-32G) and coated overnight with 1 mg/mL PDL in borate buffer. Live cell experiments utilized 35 mM Cellvis dishes. For these microscopy-based assays, 400,000 hippocampal neurons were seeded onto coverslips and fed at DIV3 and DIV7 as described above. Cells were fixed by adding warm 4% PFA in 2x PHEM buffer (120 mM PIPES, 50 mM HEPES, 20 mM EGTA, 4 mM MgSO4, and 0.24 M sucrose) directly to the media for 20 min at room temperature. Coverslips were washed 3x with PBS. After fixation, cells were permeabilized with 0.1% TritonX-100 for 10 minutes and blocked with 10% donkey serum (Fisher Scientific Cat #OB003001) or 3% BSA for 30 min. Cells were incubated for 1 hour (RT) in a primary antibody solution (in 1% donkey serum or BSA in PBS), wash 3 times (5 minutes each), incubated with secondary antibody solution (in 1% donkey serum or BSA in PBS) for 1 hr at room temperature, and washed again 3 times (5 minutes). Coverslips were mounted in Tris/glycerol/n-propyl-gallate–based mounting media and sealed with nail polish.

### Microscope Descriptions

All widefield imaging was performed using an inverted microscope (IX83-ZDC2; Evident/Olympus) equipped with a cMOS camera (Orca-Fusion, Hamamatsu) and Xenon light source. Images were acquired using a 100× 1.59 NA TIRF objective (Olympus) and with Cellsens software (Evident/Olympus).

Confocal microscopy was performed on an inverted laser scanning confocal microscopy (LSM780 or 980, Zeiss). The LSM780 was equipped with a motorized X,Y stage, Z-focus, five lasers (405, 488, 561, 594, 633), and three fluorescence detectors (two flanking PMTs and a central 34 channel GaGasp with 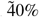 QE). Images were acquired with a 63x Plan-Apochromat oil objective (NA=1.4) using ZEN Black v 2.3. The LSM980 is equipped with a motorized X,Y stage and Z focus with high speed Piezo insert as well as four diode lasers (405, 488, 561, 633). Images were acquired with 1x Nyquist sampling using a 63x 1.4 NA Plan-Apochromat oil objective (Zeiss) and 4 channel GaAsP detectors on Zen Blue 3.6 software. This microscope is equipped with Airyscan 2, however, this module was not used in these experiments.

### Image Analysis

For quantification of dendritic filopodia and spine number, maximum projections of GFP-CAAX transfected hippocampal neurons were created from confocal z-stacks. Filopodia and spine number were manually quantified along secondary/tertiary branches off the primary dendrite of pyramidal neurons and normalized to the total length of dendrites analyzed. In general, all secondary/tertiary dendrites clearly visible (not obscured by a crossing axon or other dendrite branch) in the image were used for analysis) with a minimum of 100 μm of dendrite analyzed. Spine length and width were manually measured in FIJI. Images were blinded before analysis.

Quantification of synaptic marker puncta was completed in Imaris. Confocal z-stacks were imported into Imaris. Neurons were masked using the MAP2-AF405 channel. The remaining three channels (Synaptophysin-AF647, Gephyrin-AF594, and PSD-95-AF488) were masked and duplicated for analysis—removing any puncta outside of the MAP2 mask prior to analysis. Puncta were each marker were identified using the Imaris spots function—using 0.5 μm estimated diameter for Gephyrin and PSD-95 and 0.75 μm diameter for Synaptophysin. The colocalize spots function was then used to analyze colocalization (within 0.5 μm of spot center) between Synaptophysin/Gephyrin and Synaptophysin/PSD95. These colocalized puncta were considered inhibitory and excitatory synapses, respectively. The mean fluorescence intensity for colocalized puncta (of each protein) was exported and plotted.

Quantification of phalloidin and ArpC2 levels in dendritic spines was completed manually in FIJI. Stubby/mushroom spines were identified by eye using the phalloidin channel. These spines were manually traced and the mean fluorescence for the phalloidin and ArpC2 channels was measured for each spine. Background subtraction was completed by measuring 3 different 3 μm x 3 μm regions outside the neuron, averaging these values for each channel, and manually subtracting these values from the measured spine values. Images were blinded before analysis.

### Biotinylation Assays

For biotinylation assays, 750,000 cortical neurons were plated into each well of a six-well culture dish and fed at DIV 3 and DIV 7 as described above. On DIV 14, neurons were treated with 200 μg/mL of netrin-1 or sham media for 45 minutes. All of the following steps occurred on ice or at 4°C. Cells were washed twice with cold PBS-CM (PBS containing 1 mM MgCl2 and 0.1 mM CaCl2, pH 8.0) and incubated with 0.5mg/mL Sulpho-NHS-SS-Biotin (Thermo Scientific) for 30 minutes. Cells were briefly washed once with PBS-CM, quenched with two 20 mM glycine in PBS-CM washes (5 minutes each), and washed again with PBS-CM. After lysis, cells were rotated for 30 minutes and cleared at 14,000 rpm (sub x g equivalent). 100 μg of lysate was incubated with Neutravidin agarose beads (Thermo) for four hours. Beads were washed four times in lysis buffer and proteins were eluted with 2x SB containing BME (65°C, 15 minutes).

### Western Blotting

Samples were loaded on a 7.5% SDS-PAGE gel, transferred on nitrocellulose membrane for 90 min (75 V), blocked in 5% milk or 5% BSA in TBST for 1 hr (RT), and incubated with primary antibody overnight in 1% milk in TBST. Membranes were washed 3x (5 minutes) in TBST, incubated with secondary antibodies for 1 hr, washed 3x (10 minutes) in TBST, rinsed in PBS and imaged on an Odyssey (LI-COR Biosciences).

### Subcellular Fractionation

Following genotyping to identify *Trim9*^*+/+*^ and *Trim9*^-/-^ littermates, three week old mice were sacrificed. The forebrain (cortex and hippocampus) was dissected out in ice-cold dissection media, transferred to an Eppendorf and flash frozen in liquid nitrogen. Tissue was stored at –80C until use.

All subsequent steps were performed at 4°C. Tissue was homogenized in ice cold homogenization buffer (10 mM HEPES pH 7.4, 320 mM sucrose, 1 mM EDTA, 5 mM sodium pyrophosphate, 1 mM sodium vanadate, 150 μg/mL PMSF, 2 μg/mL leupeptin, 2 μg/mL aprotinin, and 50 μM PR-619) and dounced 15x. After centrifuging at 800 x g (10 minutes) to remove the nuclear fraction, the supernatant was transferred and centrifuged at 16,000 *x g*. This pellet was resuspended in fresh homogenization buffer and loaded onto a discontinuous sucrose gradient containing 1.2 M, 1 M or 0.8 M sucrose, as well as the inhibitors listed above. Following centrifugation in a SW-41 rotor at 82.5 k x g (90 minutes), the interface between the 1.2 and 1 M sucrose (synaptosome fraction) was collected. This interface was diluted with buffer to return the sucrose concentration to 320 mM and centrifuged at 100k *x g* (30 minutes). This pellet was resuspended in 50mM HEPES buffer (plus inhibitors) and then mixed at a 1:1 ratio with 50mM HEPES buffer with 1% Triton (Final concentration = 0.5% Triton). Following a 15 minute rotating incubation, the samples were centrifuged at 32k *x g* (20 minutes). The pellet (post-synaptic density fraction) was re-suspended in 50 mM HEPES and flash frozen in liquid nitrogen.

Following protein concentration quantification by Bradford, 100 μg of protein was aliquoted from each PSD. Two PSD samples of each condition were combined and a total of four replicates (from eight mice) for each condition were submitted to the UNC Hooker Proteomics Core.

### Proteomics

Lysates (0.2 mg per sample; 4 replicates per condition) were precipitated using 4x cold acetone and stored at -20ºC overnight. The next day, samples were centrifuged at 15000xg at 4ºC for 15 min, then protein pellets were reconstituted in 8M urea. All samples were reduced with 5mM DTT for 45 min at 37ºC, alkylated with 15mM iodoacetamide for 30 min in the dark at room temperature, then diluted to 1M urea with 50mM ammonium bicarbonate (pH 7.8). Samples were digested with LysC (Wako, 1:50 w/w) for 2 hr at 37ºC, then digested with trypsin (Promega, 1:50 w/w) overnight at 37ºC. The resulting peptide samples were acidified, desalted using desalting spin columns (Thermo), then the eluates were dried via vacuum centrifugation. Peptide concentration was determined using Quantitative Colorimetric Peptide Assay (Pierce).

A total of 16 samples (50 μg each) were labeled with TMTpro reagents (Thermo Fisher) for 1 hr at room temperature. Prior to quenching, the labeling efficiency was evaluated by LC-MS/MS analysis. After confirming >98% efficiency, samples were quenched with 50% hydroxylamine to a final concentration of 0.4%. Labeled peptide samples were combined 1:1, desalted using Thermo desalting spin column, and dried via vacuum centrifugation. The dried TMT-labeled sample was fractionated using high pH reversed phase HPLC (Mertins *et al*., 2018). Briefly, the samples were offline fractionated over a 90 min run, into 96 fractions by high pH reverse-phase HPLC (Agilent 1260) using an Agilent Zorbax 300 Extend-C18 column (3.5-μm, 4.6 × 250 mm) with mobile phase A containing 4.5 mM ammonium formate (pH 10) in 2% (vol/vol) LC-MS grade acetonitrile, and mobile phase B containing 4.5 mM ammonium formate (pH 10) in 90% (vol/vol) LC-MS grade acetonitrile. The 96 resulting fractions were then concatenated in a non-continuous manner into 24 fractions and 5% of each were aliquoted, dried down via vacuum centrifugation and stored at -80ºC until further analysis.

The 24 TMT labeled proteome fractions were analyzed by LC/MS/MS using an Easy nLC 1200-Orbitrap Fusion Lumos (Thermo Scientific). Samples were injected onto an Easy Spray PepMap C18 column (75 μm id × 25 cm, 2 μm particle size) (Thermo Scientific) and separated over a 120 min method. The gradient for separation consisted of 5–42% mobile phase B at a 250 nl/min flow rate, where mobile phase A was 0.1% formic acid in water and mobile phase B consisted of 0.1% formic acid in 80% ACN. The Lumos was operated in SPS-MS3 mode (McAlister *et al*., 2014), with a 3s cycle time. Resolution for the precursor scan (m/z 400–1500) was set to 120,000 with a AGC target set to standard and a maximum injection time of 50 ms. MS2 scans consisted of CID normalized collision energy (NCE) 32; AGC target set to standard; maximum injection time of 50 ms; isolation window of 0.7 Da. Following MS2 acquisition, MS3 spectra were collected in SPS mode (10 scans per outcome); HCD set to 55; resolution set to 50,000; scan range set to 100-500; AGC target set to 200% with a 100 ms maximum inject time.

Raw data files were processed using Proteome Discoverer version 2.5, set to ‘reporter ion MS3’ with ‘16pex TMT’. Peak lists were searched against a reviewed Uniprot mouse database (downloaded Feb 2021 containing 17,051 sequences), appended with a common contaminants database, using Sequest HT within Proteome Discoverer. Data were searched with up to two missed trypsin cleavage sites, fixed modifications: TMT16plex peptide N-terminus and Lys, carbamidomethylation Cys, dynamic modification: N-terminal protein acetyl, oxidation Met. Precursor mass tolerance of 10ppm and fragment mass tolerance of 0.5 Da (MS3). Peptide false discovery rate was set to 1%. Reporter abundance was calculated based on intensity; for MS3 data, SPS mass matches threshold was set to 50 and co-isolation threshold was set to 100. Razor and unique peptides were used for quantitation. Proteins with >50% missing TMT intensities across samples were removed. Student’s t-tests were conducted within Proteome Discoverer, and a p-value <0.05 was considered significant. Log_2_fold change ratios were calculated for each pairwise comparison.

Protein-protein interactions were visualized with Cytoscape (Shannon *et al*., 2003). Enrichment of biological processes in significantly changed proteins was identified using Gene Ontology analysis (Ashburner *et al*., 2000; Thomas *et al*., 2022; The Gene Ontology Consortium *et al*., 2023).

## ACKNOWLEDGEMENTS

We thank Vong Thoong, Christopher Hardie, Mayra Correa-Ramirez, Janee CadlettJette, and Natallia Riddick for assistance with mouse husbandry and genotyping. We thank Blake Creighton, Sean Gay, and Michael Ye for technical troubleshooting and helpful discussions over the years, and Katie Baldwin for immunofluorescence advice. This work was supported by National Institutes of Health Grants R35GM135160 (S.L.G.), 1F31NS113381-01 (L.E.M). Mass spectrometry was performed at the UNC Proteomics Core Facility, which is supported in part by NCI Center Core Support Grant (2P30CA016086-45) to the UNC Lineberger Comprehensive Cancer Center. Mass spectrometry was also supported by an award from the UNC Core Facilities Advocacy Committee and Office of Research Technologies, UNC Chapel Hill School of Medicine. Confocal microscopy was performed at the UNC Neuroscience Microscopy Core, which receives funding from the NIH-NINDS Neuroscience Center Support Grant P30 NS045892 and the NIH-NICHD Intellectual and Developmental Disabilities Research Center Support Grant P50 HD103573. In particular, this work utilized the LSM980 microscope at the microscopy core, which was funded with support from NIH grant S10 OD032388.

