## Supplemental Material for "The E3 ubiquitin ligase TRIM9 regulates synaptic function and actin dynamics"

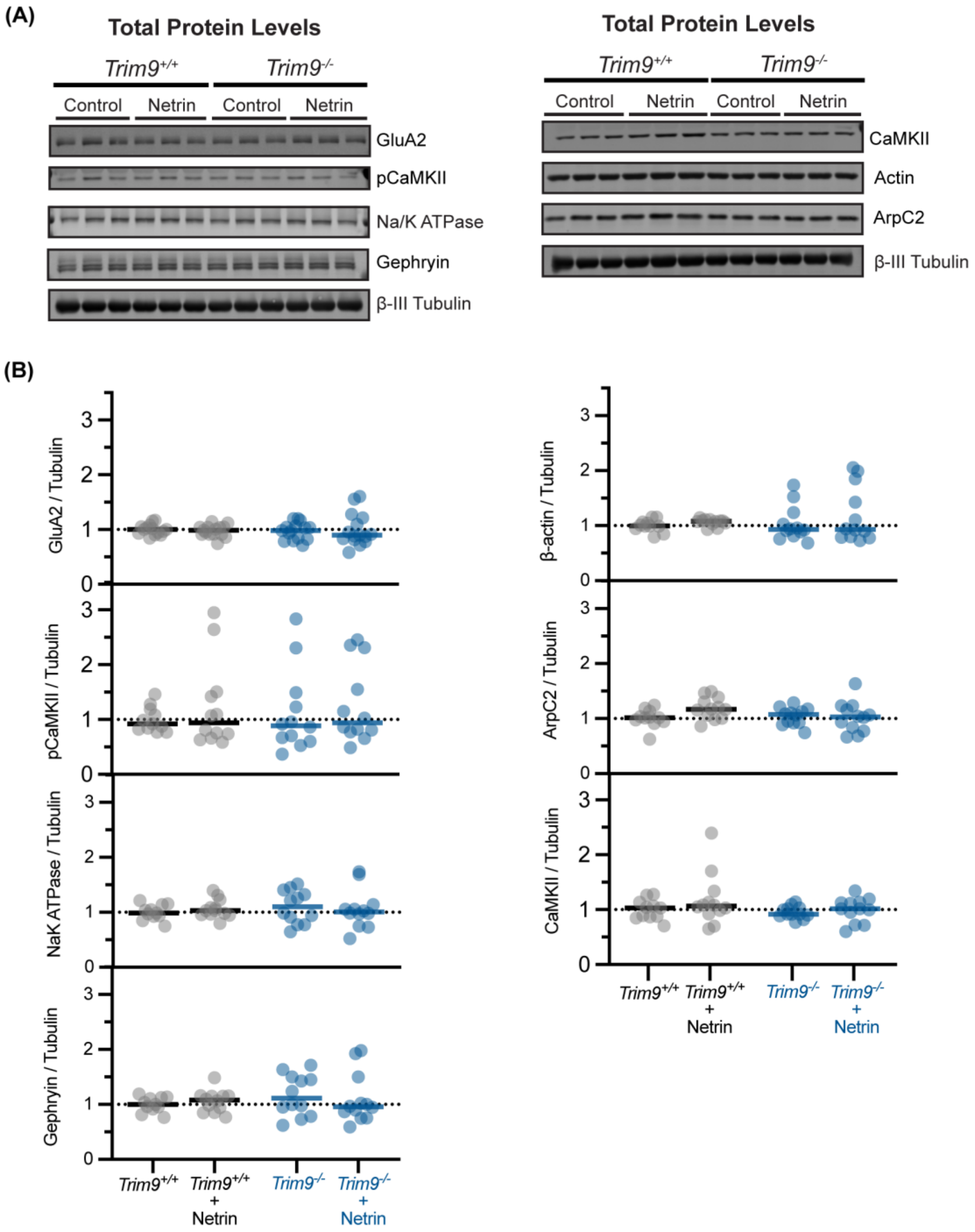


**Figure S1. Loss of *Trim9* does not alter numerous dendritic spine components.** Quantitative western blotting of total protein levels from cultured cortical neurons (DIV 14), treated with netrin-1 (200 µg/mL) or sham media for 45 minutes. N = 11-12 technical replicates across independent experiments. A Kruskal-Wallis test with Dunn's multiple comparisons was completed for each protein.
